# Cas12a cleavage and trimming kinetics reveal mismatches as a strategy to steer editing

**DOI:** 10.64898/2026.08.31.748204

**Authors:** Uzair Ahmed, Fausta Michnevičiūtė, Marius Vinogradovas, Eimina Dirvelytė-Valauskė, Urtė Neniškytė, Stephen Knox Jones

## Abstract

Gene knockouts by CRISPR-Cas nucleases rely on targeted DNA cleavage and error-prone DNA repair: end-joining pathways can introduce insertions and deletions that assist in disrupting the coding sequence. However, only a fraction of edits achieves this, and an unfavorable array of repair outcomes typically requires switching to another editing technology. Key factors that influence repair are the types and lengths of DNA ends following cleavage. Here, we investigated Cas12a’s ability to produce different ends and if they can be used to redistribute editing outcomes.

We determined the sites and rates of target cleavage by Cas12a *in vitro* by combining kinetic modeling with nucleotide-resolution assays. Here, we show that trimming – repeated cleavage of an already cut target – occurs about 4x faster than initial cleavage; it also presents alternative DNA end structures for cellular repair. We also introduced specific mismatches to the gRNA: Cas12a maintained fast target cleavage, but changed where the target was cleaved and how quickly it was trimmed, compared to matched gRNA.

We exploited the differences in cleavage kinetics between matched and mismatched gRNAs to develop reprogrammed gRNAs, *i.e.* rpgRNAs. Intentionally-mismatched rpgRNAs retained the high editing efficiency observed with traditional gRNAs. However, they shifted editing between in-frame and out-of-frame outcomes to enhance gene knockout success across genes. Reprogrammed gRNAs offer an efficient way to steer editing toward such preferred outcomes, while retaining the simplicity of gene editing with CRISPR-Cas nucleases.

## Introduction

CRISPR-Cas nucleases underly this century’s transition to targeted, precise and robust genome editing technologies ^1^. In the simplest case, we employ nucleases – such as Cas9 and Cas12a – across organisms and cell types for efficient gene knock out^2^. Guide RNAs (gRNAs) direct a Cas nuclease to its target site in the genome. This RNA programmability reduces the time and cost of gene editing, helping overcome global challenges in research, chronic disease, and food security ^3,4^. But for successful cleavage, the target must be positioned beside a short protospacer adjacent motif (PAM) ^5^. Once identified, the nuclease cleaves its target, producing a double-stranded break (DSB) in the DNA ^6^.

End joining pathways like non-homologous end joining (NHEJ) repair DSBs at the target, often inadvertently introducing insertions or deletions (indels) in the target site sequence ^7^. An indel edit can cause a frameshift in protein-coding genes; the subsequent loss of function marks a successful gene knock-out ^8,9^. However, indel edits of length 3N (*e.g.* -6, -3, 0, or 3 bps) instead cause codon loss, codon gain, or mutations. Here, continued function, even if attenuated, marks an unsuccessful or partial knock-out ^10^.

The editing outcome for a cell population contains an array of edits, depending on the site targeted, the cell type, and the nuclease used^11^. But users cannot control which edits occur, or how frequently; even a highly efficient Cas nuclease may not yield knock-out success. Repeating an editing experiment produces nearly the same array of edit outcomes, and thus cannot improve success rates^12,13^. While repair pathway modulators or different technologies (like prime editing ^14^) can help, this remains a technological gap for gene knock outs with Cas nucleases.

We, and others, have proposed that the type of ends produced by Cas nucleases may alter the outcomes of DSB repair ^15,16^. With *Sp*Cas9, the engineered “LZ3” variant changes the frequency of cutting targets to generate +1 overhang ends (compared to natural *Sp*Cas9) and contribute to +1 insertion frameshift edits ^15,17^. Meanwhile, *As*Cas12a generates larger overhangs than *Sp*Cas9; their position and lengths depend on the target site, gRNA-target pairing and exposure time ^16,18^. Further, *AsCas12a* repeatedly cuts (*i.e.* trims) exposed ends after initially cutting its target ^19^. Full removal of trimmed nucleotides prevents base-pairing that would otherwise aid in scarless repair. Despite this, trimming has never been employed as a strategy for altering editing outcomes.

Here, we address these gaps by characterizing *As*Cas12a trimming kinetics and then pursuing trimming differences due to gRNA-target mispairing as a strategy to alter editing outcomes. By combining nucleotide-resolution biochemistry and mathematical modeling, we find that the ends produced by Cas12a can result from trimming rather than initial cleavage, and that these processes occur on similar timescales. Cas12a programmed with gRNAs containing cleavage-site adjacent mispairs showed a similar overall rate of cleavage to gRNAs without mispairs. However, mispairs shifted the sites of DNA cleavage and the rates of cleavage for each strand and position. We then performed editing experiments with these gRNAs: gRNAs programmed for intentional mispairing – *i.e.* reprogrammed gRNAs (rpgRNAs) – retain high editing efficiency, but produced significantly different editing outcomes, shifting the frequency of codon loss vs frameshift mutations during knock-out experiments.

Together, these findings represent the first quantitative measures of trimming rates for any CRISPR-Cas nuclease, and they establish mispairing via rpgRNAs as an effective strategy to increase favorable repair outcomes in human cells while retaining the simplicity of standard CRISPR-Cas nucleases.

## Results

### Cas12a retains fast cleavage of off-targets with PAM-distal mismatches, but cleaves them differently

We set out to systematically map Cas12a’s cleavage kinetics and cut site differences with mismatched target-gRNA pairs (Figure 1, A-F). Based on our previous high-throughput *in vitro* data, we selected targets with cleavage-site adjacent mismatches (position 19 from the PAM: MM19; Figure 1B), as Cas12a cleaves such targets at similar rates to an on-target, but with different cleavage patterns ^16^. To observe target cleavage with single-nucleotide resolution, we tracked individual DNA ends and strands over time with denaturing PAGE (Figure 1C, Supplemental Figure S1). Incubating Cas12a with linear DNAs (with offset targets and end labels; Figure 1A) allowed us to visualize changes in the distribution of each strand’s PAM-proximal and PAM-distal products (hereafter referred to as distal and proximal, or P and D) ^20^. This provided unique insight into how mismatches force Cas12a to deviate from its canonical cleavage positions.

**Figure 1.**
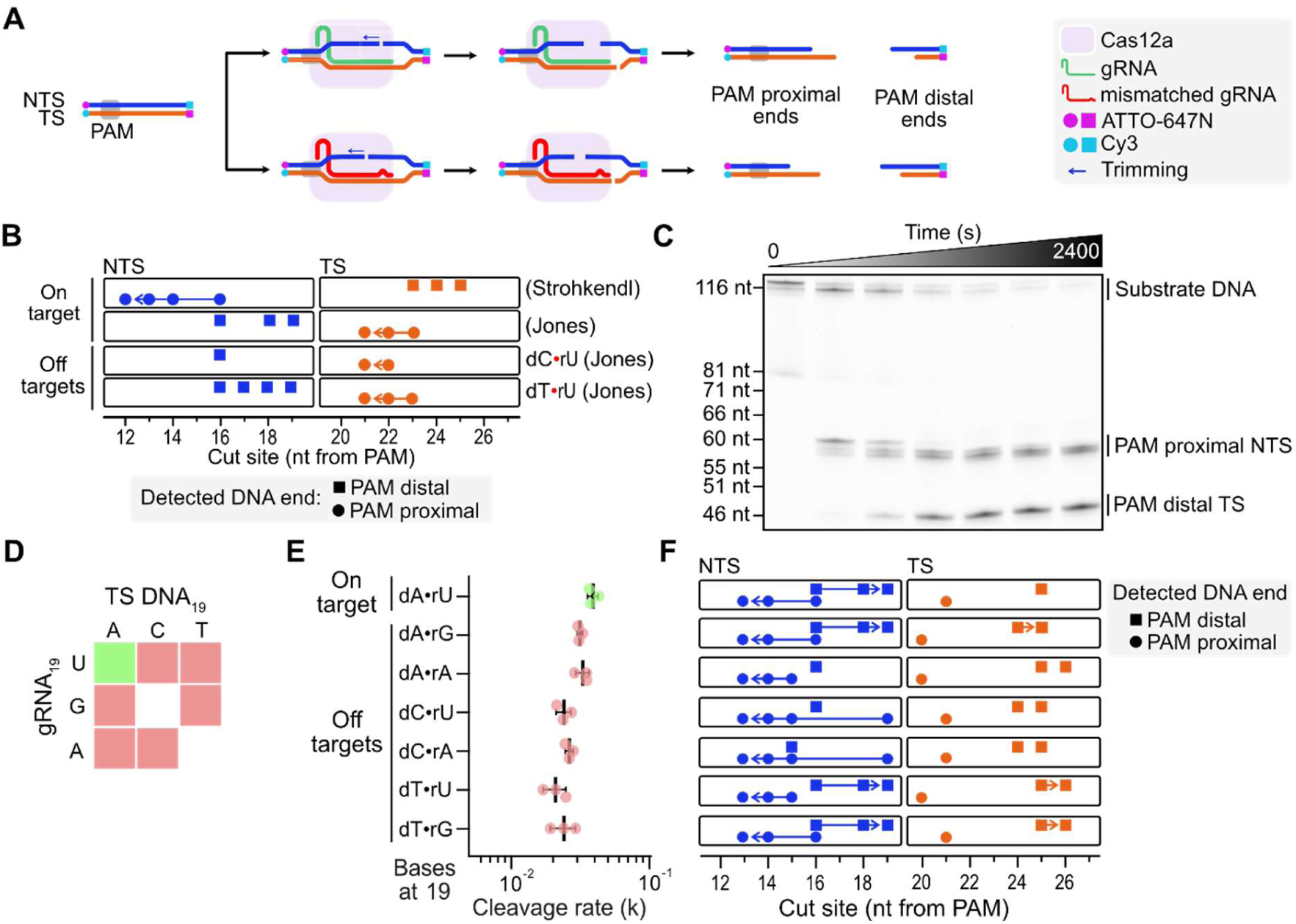
Cas12a cleavage sites depend on gRNA-target pairing. **A)** Schematic of Cas12a cleavage of on-targets (upper) and off-targets (lower). Target DNAs contained fluorophores on all ends (5’ ends: ATTO-647N, 3’ ends: Cy3). **B)** Previously-reported cut sites for on-target PAM-proximal NTS and PAM-distal TS strands (Strohkendl) ^18^, and for on- and off-target PAM-distal NTS and PAM-proximal TS strands (Jones) ^16^. Arrows indicate trimming. Cut sites are shown as nucleotides (nt) from the PAM. **C)** Single nucleotide resolution denaturing urea-PAGE showing products formed from on-target DNA cleavage (ATTO-647N 5’-tagged PAM-proximal NTS and PAM-distal TS products shown). **D)** Target DNAs tested in this study. Off-targets contained the indicated TS DNA-gRNA mismatches at position 19. Green: on-target, red: off-targets. **E)** Cleavage rates for on-target and off-targets from D. Green: on-target, red: off-targets. N=3. Mean ± SD. **F)** Cut sites for all on-target and off-target DNA strands. Arrows indicate trimming. Cut sites are shown as nucleotides (nt) from the PAM.

To determine if different mismatch identities alter Cas12a cleavage activity, we tested Cas12a with all possible target DNA and gRNA mismatches at position 19 (Figure 1D). The measured cleavage rates of all mismatch combinations were similar to the on-target cleavage rate (0.02 to 0.03 s^-1^; on-target k= 0.03 s^-1^; Figure 1E).

We then compared the Cas12a cut sites that we observed for these targets with those observed in prior NGS-based ^16^ and gel-based studies ^18,20^ (Figure 1B,F; Supp. Figure 1). For an on-target, the NGS-based study reported proximal target strand (TS) cut sites at positions 21-23 with repeated time-dependent cleavage, *i.e.* trimming, from position 23 down to 21. Distal non-target strand (NTS) cuts occurred at positions 16-19, but NTS trimming was undetectable due to end repair in NGS sample preparation. The gel-based study reported cut sites at distal TS positions 23-25 but could not detect TS trimming due to 5’ radiolabeling. Proximal non-target strand (NTS) cut sites were at positions 12-16, with possible trimming from position 16 down to 12.

Our data connects these disparate results by reporting cleavage sites for all four DNA ends from both strands (Figure 1F; as in Cofsky *et al* ^21^ and Wörle *et al* ^20^). We found that Cas12a cleaves and trims the on-target NTS at multiple sites from position 19 to 13, while the TS was cleaved at position 21 and 25 (our assay does not distinguish ssDNA and dsDNA cleavage events).

The prior NGS-based study also reported cut sites for different MM19 targets, where the off-target NTSs were cleaved from position 19-16 and TSs were cleaved from position 24-21 ^16^ (only distal NTS and proximal TS ends were reported). We detected Cas12a cleavage of these off-targets at an even wider range of positions: Off-target NTS can be cleaved from position 19-13 with trimming, whereas off-target TS can be cleaved from position 26-20, depending upon the mismatch (Figure 1F).

With our design, we mapped the cleavage products from each DNA end, yielding a comprehensive view of how Cas12a cleaves these on- and off-targets. The identities of the mismatched nucleotides at position 19 result in different cleavage profiles – presumably, mismatched bases distort the DNA, shifting the sites of cleavage. As cleavage also requires multiple Cas12a domains to undergo conformational changes, mismatches further modulate these dynamics ^19,21,22^.

### An ODE system captures dynamics of DNA trimming

Prior experiments qualitatively report on products of Cas12a cleavage and its ability to trim them, but do not report kinetic rates for trimming ^16,18,20^. How does Cas12a operate on products already associated with it, positioned very near to its RuvC nuclease domain? The null hypothesis assumes that Cas12a does not distinguish between intact and already-cleaved DNAs, such that the rate of cleavage to a product should not differ whether from an intact “substrate” DNA or from a longer “product” DNA. Yet, trans-cleavage by Cas12a nucleases indicates that initial cleavage events alter subsequent cleavage events ^18^. Thus, we next investigated how Cas12a operates on products already associated with it.

To capture the dynamics of cleavage, we developed a system of ordinary differential equations (ODEs) that fits initial product formation and subsequent trimming rates with first-order kinetics under saturating enzyme conditions. While designed for Cas12a, the system generalizes to any nuclease with any number of initial cleavage sites or trimming possibilities (https://github.com/JonesLabEU/gRNA-reprogramming). The system requires the normalized intact substrate target DNA fraction (*S*), and proximal and distal DNA cleavage product fractions (*P_Pn_, P_Dn_*, where n = product number) for each time (*t*). The system fits initial product formation rates directly from the substrate (*k_1_, k_2_,… k_n_*) and subsequent products formation rates due to trimming (*k_1-2_, k_2-3_, …, k_n-n+1_, k_1-3_, …, k_n-n+2_, etc*.). Input data are separated into sets based on their labels: 5’ tagged ends (proximal NTS and distal TS) or 3’ tagged ends (distal NTS and proximal TS) and modeled with ODE systems 1 and 2, respectively (Figure 2A).

**Figure 2.**
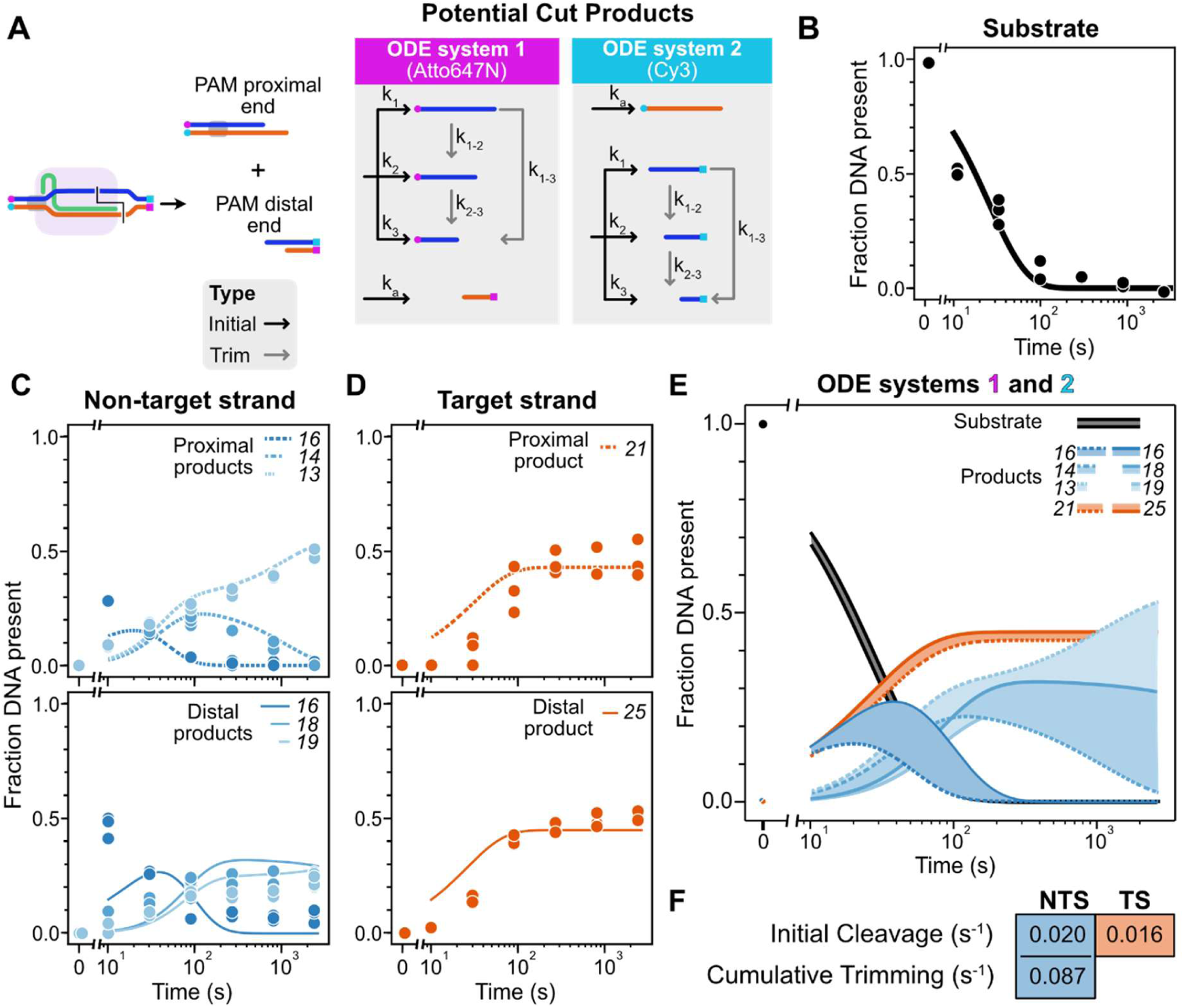
Systems of ordinary differential equations capture the dynamics of on-target DNA cleavage and trimming by Cas12a. **A)** Schematic models of the ODE systems. Cas12a cleaves the target DNA into PAM-proximal and PAM-distal ends with a staggered cut. The ODE systems detect the rates of formation for all products and the rates of any subsequent trimming. **B)** On-target substrate DNA cleavage by Cas12a. Curve: Model-fitted depletion of on-target DNA (measured from 5’ tags). N=3. **C)** On-target NTS products generated by Cas12a. Blue shades: products of cleavage at the indicated distance from the PAM. Curves: Model-fitted fraction of the indicated product (measured from 5’ and 3’ tags). **D)** On-target TS products generated by Cas12a. Orange shades: products from cleavage at the indicated distance from the PAM. Curves: Model-fitted fraction of the indicated product (measured from 5’ and 3’ tags). Rates for C and D generated from the mean of three independently-fitted biological datasets. **E)** Complete modelled fractions of Cas12a substrates and products for all DNA ends. Substrate depletion rates are shown in black, NTS products in shades of blue and TS in orange. Thick lines: PAM-proximal ends, thin lines: PAM-distal ends. Fills assist with identifying product curves. **F)** Model-fitted initial cleavage and cumulative trimming rates for each strand.

We used the ODEs to model Cas12a activity with an on-target DNA substrate (Figure 2). The overall substrate cleavage rate of 0.03 s^-1^ agreed with prior studies ^16,18^ (Figure 2B). Analyzing the three proximal NTS products, we found that Cas12a initially only generated the longest product (*P_P16_*,), but rapidly trimmed it by 2-3 nucleotides into shorter products (*P_P14_, P_P13_*), until only *P_P13_* remained (0.087 s^-1^, ODE system 1, Figure 2C). Cas12a similarly generated three distal NTS products: Cas12a initially only made *P_D16_*, but quickly trimmed it into the shorter products *P_D18_, P_D19_* (0.023 s^-1^, ODE system 2, Figure 2C). This indicated that when Cas12a cuts on-target DNA, it favors trimming ends from already-associated products rather than from the substrate. Analyzing the TS, we found that Cas12a generated only single products (*P_D25_* and *P_P21_*, 0.016 and 0.020 s^-1^) without subsequent trimming (Figure 2D). The positional difference between TS product ends may represent trimming that occurs faster than can be detected (trim *k* >>> initial cleavage *k*).

Between the two ODEs, we mapped the kinetics of all DNA ends produced by Cas12a cleaving an on-target DNA (Figure 2E). We found that while Cas12a generated single TS products, cumulatively, NTS trimming events occur ∼4x faster than initial target cleavage (Figure 2F).

### Cas12a cleaves DNA ends in a target-specific manner

Mismatches between gRNAs and targets – especially near cleavage sites – can dramatically alter cleavage kinetics for Cas12a and other Cas nucleases ^16,18^. For Cas9, this repositions DNA strands relative to its nuclease domains (as captured with recent structures) and depends on the specific mispair formed^23,24^. To understand how gRNA-DNA mismatches near potential cleavage sites impact Cas12a activity, we investigated off-target cleavage and trimming using gRNAs and target DNAs that form mismatches at the 19^th^ position from the PAM (MM19; Figure 1D).

We found that Cas12a operates on an MM19 off-target (dC▪rA) to generate different products at different rates than for an on-target (Figure 3, other off-targets: Supplemental Figure S2). Cas12a initially cleaved off-target DNA substrate into four different proximal NTS products (*P_P19_*, *P_P15_*, *P_P14_* and *P_P13_*), rather than one as for on-target (*P_P15_*, Figure 3A vs C). Still, position 15 was the preferred cleavage site, with a 10x or faster rate (0.016 s^-1^) than for the others (≤0.0016 s^-1^). Cas12a still trimmed all the products to the shortest one (*P_P13_*; Figure 3B). Conversely, Cas12a generated a single distal NTS product (*P_P15_*), but further trimming was abolished. Finally, Cas12a generated a single proximal TS product (*P_P21_*), but generated two distal TS products (*P_D24_* and *P_D25_*) at similar rates that also were not trimmed (Figure 3C). We reported the differences in cleavage and trimming rates for other mismatched targets similarly (Figure S2, Supp. file 2).

**Figure 3.**
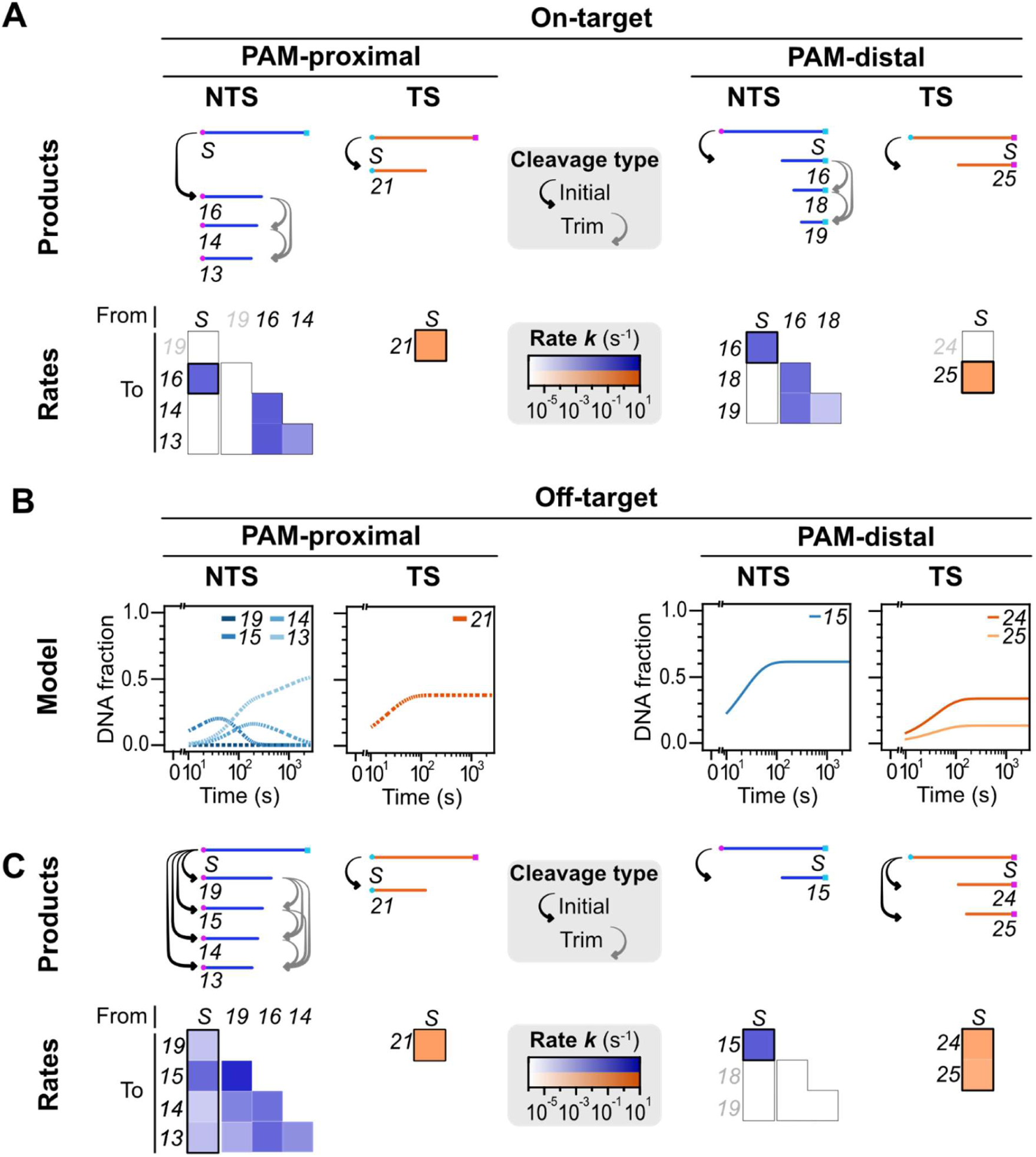
Kinetics of Cas12a cleavage and trimming of an on-target and off-target. **A)** Products formed from initial cleavage (black arrows) and trimming (grey arrows) of the on-target. Model-fitted rates of conversion from substrates (S) to products, or from one product to another. Numbers identify products by the distance from their cleavage site to the PAM. Blue shades: log-scale rates for the NTS. Orange shades: log-scale rates for the TS. Bolded boxes: products initially cleaved from the substrate. **B)** Model-fitted fractions of cleavage products for an off-target (dC•rA), generated by Cas12a. Blue shades: NTS products at the indicated distance from the PAM. Orange shades: TS products at the indicated distance from the PAM. Thick lines: PAM-proximal ends, thin lines: PAM-distal ends. Rates generated from the mean of three independently-fitted biological datasets. **C)** Products formed from initial cleavage (black arrows) and trimming (grey arrows) of an off-target (dC•rA). Model-fitted rates of conversion from substrates (S) to products, or from one product to another. Numbers identify products by the distance from their cleavage site to the PAM. Blue shades: log-scale rates for the NTS. Orange shades: log-scale rates for the TS. Bolded boxes: products initially cleaved from the substrate.

Collectively, we found that the lengths and total number of NTS and TS products differ when Cas12a’s gRNA and target mismatch at position 19. This mismatch could completely prevent Cas12a from trimming the distal NTS, even when it trimmed other ends similarly across the targets. Given that Cas12a processed DNA ends in a target specific manner, we next investigated if reprogramming Cas12a with a single, intentionally-mismatching (MM19) gRNA could alter editing outcomes.

### Reprogrammed gRNAs produce different editing outcomes

We hypothesized that mismatches can be used to generate different editing outcomes with Cas12a without reducing overall editing efficiency. We programmed Cas12a with either fully-matched control gRNAs or gRNAs with a target-relative mismatch at position 19, which we refer to as reprogrammed guide RNAs (rpgRNAs). Such RNPs were electroporated into human cells, and after three days, we sequenced the target site to determine editing efficiency and map editing profiles ^25^ (Figure 4A, 4B). Two rpgRNAs (A had a dA▪rA mismatch; B had a dA▪rG mismatch, similar to Figure 1F) targeting *CD96* in NK-92 cells maintained editing efficiencies similar to control gRNA (∼60% indel formation; Figure 4C). However, they generated a different array of editing outcomes when compared to the control gRNA.

**Figure 4.**
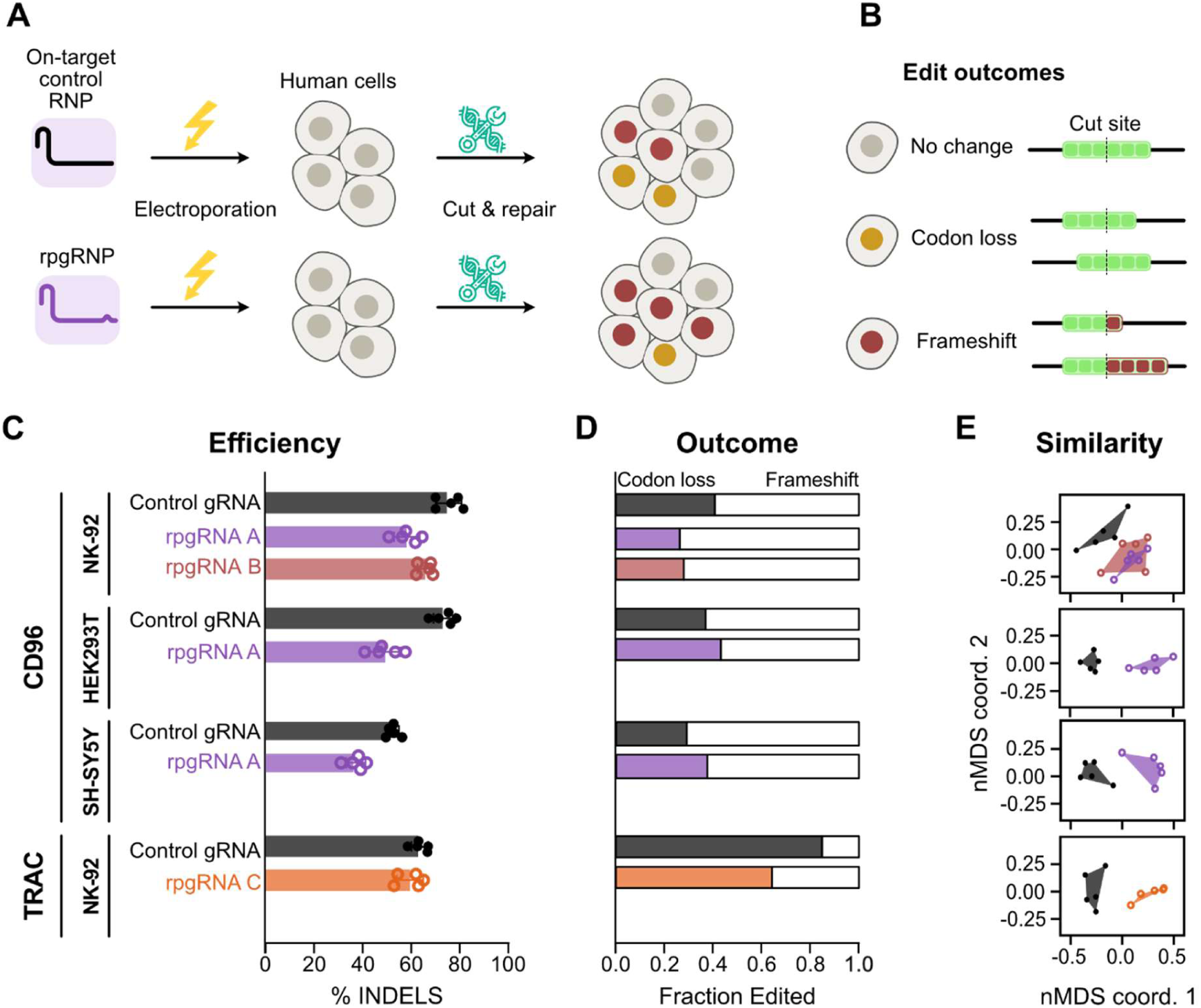
Reprogrammed gRNAs (rpgRNAs) alter repair outcomes in human cells. **A)** Schematic of rpgRNA usage. Cas12a RNPs containing either gRNAs or rpgRNAs are electroporated into human cells. After targeted cleavage, cellular DNA repair can produce different editing outcomes depending on the RNP used. **B)** Editing outcomes can include no change, codon loss (or gain), or a frameshift in the target gene. **C)** Cas12a editing efficiency with gRNAs and rpgRNAs. Genes and cell types are indicated on the left. Colors: different gRNAs. N=5. Mean ± SD. **D)** Editing outcomes with gRNAs or rpgRNAs across different genes and cell types. Colored bars: codon loss, white bars: frameshift. **E)** Similarity of editing outcomes with gRNAs or rpgRNAs across different genes and cell types by nMDS analysis. N=5.

The editing outcomes (almost always deletions; Supp. file 3) from rpgRNAs A and B in NK-92 cells were more often frameshifts compared to control gRNA (73% vs 59%, Figure 4D). While codon loss occasionally results in loss-of-function, frameshift mutations almost always do. To determine the significance of this shift, we performed nMDS analysis (Figure 4E). The rpgRNAs and control gRNA generated significantly different arrays of edit outcomes for the CD96 locus (N=5, Shepard plot stress value: 0.15, PERMANOVA: p=0.039, Figure 4E), while outcomes for rpgRNA A and B were similar (PERMANOVA p=0.391).

To determine the robustness of rpgRNAs for providing alternative editing outcomes, we performed similar experiments at the CD96 locus in additional cell lines and another locus. Control gRNAs and rpgRNAs achieved similar editing efficiencies (Figure 4C) but different edit outcome populations (Figure 4E; Shepard plot stress and PERMANOVA p values for CD96 in HEK293T: 0.015, p=0.0065, CD96 in SH-SY5Y: 0.057, p=0.0065, TRAC in NK-92: 0.039, p=0.0065). The editing outcome analysis again showed differences in frameshift frequencies with each rpgRNA as compared to control gRNAs, though not always in the same direction (Figure 4D; CD96 in HEK293T: 57% vs 64%, CD96 in SH-SY5Y: 62% vs 70%, TRAC in NK92 =36% vs 15%). Our data show that in each case and cell type tested, rpgRNAs are sufficient to change repair outcomes with Cas12a between codon losses and frameshifts, or vice versa.

We can reprogram DNA repair outcomes using rpgRNAs with Cas12a, which provides a simple, efficient and cost-effective solution to pivot repairs toward desired editing outcomes without losing editing efficiency.

## Discussion

This study characterizes Cas12a nuclease by assessing its cleavage kinetics and sites, and is the first to quantify its trimming kinetics. We show that Cas12a operates differently on matched and mismatched targets; it cuts them at different positions but preserves overall cleavage efficiency. Our ODE system captures nuclease cleavage and trimming kinetics, predicting how Cas12a operates on on-targets and off-targets (MM19), thus revealing that trimming occurs faster than initial target cleavage. The differences in cleavage positions upon target-gRNA mismatching prompted us to leverage intentionally mismatched rpgRNAs to alter the outcomes of gene editing with Cas12a. We show that rpgRNAs change the frequency of frameshift vs. codon loss edits and provide a new strategy for improving gene knock-out success while retaining the simple, two-component simplicity promised by Cas nucleases.

How do mismatches between gRNAs and targets induce changes in Cas12a cleavage activity and subsequent repair? One clue lies in Cas12a’s mechanism: as a Type V Cas nuclease, it relies on a single RuvC nuclease domain ^19,26^. RuvC (or Cas12a’s hold on its target) must flex to accommodate both DNA strands ^21,22^ (reviewed in ^27^); this also enables Cas12a’s trans-cleavage activity and gRNA processing ^28,29^. Mismatched bases likely distort the R-loop, and thus the position of each strand within the RuvC active site, leading to the differences observed here. However, these repositions require structural studies to truly understand.

The greater mystery lies in how differences in cleavage positions and trimming translate to different repair outcomes in cells. Repair via NHEJ can proceed with or without end resection^30^, in agreement with the indel edits that we observed (Supp. file 3). However, different overhang types and base identities play a critical role in defining how DNA is repaired, determining which strand is resolved first, and how ^31^. Beside a few exceptions with Cas9^32–34^, such editing outcomes with Cas nucleases remains unpredictable, and thus, underly the drive toward more deterministic yet complex tools, like base^35,36^ and prime editors^37^.

Beyond simply identifying cleavage products, our ODE systems reveal how fast products are made, and from which sources. They quantify initial cleavage and trimming, so we can recognize how quickly Cas12a operates on products already associated with a primed complex. However, gel and time resolution can limit the system – infrequent or extremely fast events will not be captured. Trimming activity may be rampant across Type V Cas nucleases; likely candidates include Cas12j^38^ and Cas12b^39^, which produce long overhangs (9-11 nt) that may be sensitive to mismatches^38,40,41^. Further, the general architecture of our ODE systems works for cleavage or nicking, whether by endonucleases ^42^ or exonucleases ^43^, including other Cas nucleases. Analysis of gRNA-target mispairs within Cas12a would also be valuable, as mismatches in positions 9-15 (from the PAM) bias Cas12a towards nicking – mismatches here can slow down second strand cleavage ^44^.

Rather than ‘wasting’ gRNA programming capacity, rpgRNAs repurpose their distal sequence to alter repair outcomes. This comes at almost no cost, as mismatches far from Cas12a’s PAM make almost no contribution to Cas12a’s specificity^45^, as evidenced in *in vitro* and genomic off-target analyses^46^. Thus, improved knock-out success (up to 20%; Figure 4D) can be as simple as purchasing another gRNA or performing single base mutagenesis in an encoding plasmid. However, whether frameshift mutations become more or less frequent, will depend on the gene and cell targeted. Follow-up studies should determine if rpgRNA-induced changes to editing outcomes can be predicted, similar to editing outcomes with standard gRNAs for Cas9 or Cas12a^47^. If so, rpgRNAs could also help overcome trinucleotide repeat disorders, where codons should be removed while retaining coding frame. Finally, engineered rpgRNAs may work synergistically with engineered enzymes, such as Cas12-based prime^14^ or base editors^48,49^.

## Methods

### DNA and RNA preparation and tagging

Oligonucleotides and AsCas12a gRNAs were synthesized by Genewiz (Supp. file 1). Off-target DNAs and gRNAs contained substituted nucleotides at position 19 in the spacer region. The DNA target is placed in the middle of DNA design, so when Cas12a cleaves the target PAM proximal ends run bigger on gel (Supp. figure 1). Moreover, each end has distinct fluorophore and thus can be easily distinguished on a gel. 5’ ATTO647N-tagged dsDNA targets were amplified from ssDNA oligos using 5’ ATTO-647N tagged primers (Supp. file 1) and Phusion Plus polymerase (Thermo Fisher Scientific; 30 cycles, 1X Phusion Plus reaction buffer). To tag 3’ ends of dsDNA targets, targets were treated with terminal deoxynucleotidyl transferase (TdT) (Thermo Fisher Scientific) and 7-Propargylamino-7-deaza-ddATP-Cy3 (Jena Bioscience) in 1X TdT reaction buffer (Thermo Fisher Scientific) for 2 hours at 37 °C, then quenched with 50 mM EDTA. DNA targets were purified using the GeneJET PCR purification kit (Thermo Fisher Scientific). Though we could not guarantee that all ends were labeled (see size variation in substrate DNA, Fig 1C), only tagged ends were visible and used for rate determination.

### Custom ssDNA ladder preparation and tagging

Oligonucleotides ranging from 35 to 116 nt (10 total; Supp. file 1, Genewiz) were mixed in equal mass per band ratio to form an ultra-low range (ULR) ssDNA ladder. The mix was treated with TdT and either 7-Propargylamino-7-deaza-ddATP-Cy5 or 7-Propargylamino-7-deaza-ddATP-Cy3 (Jena Bioscience) for 2 hours at 37 °C and quenched with 50 mM EDTA.

### AsCas12a production and purification

For *in-vitro* experiments, Cas12a was expressed in *E. coli* BL21(DE3) cells transformed with pET19-AsCas12a (pIF502)^18^ containing His_6_, Twin-Strep, and SUMO tags via electroporation. Cells were grown in Terrific Broth (TB) medium (24 g l−1 yeast extract, 12 g l−1 tryptone, 0.4% glycerol, 2.31 g l−1 KH2PO4, 12.54 g l−1 K2HPO4) plus carbenicillin (100 μg/mL) at 37 °C with shaking until OD600 ∼0.6. Protein expression was induced with IPTG (1mM) at 18 °C for 24 hours. Cells were harvested by centrifugation (2,500 g, 10 min, 4 °C) and stored at −80 °C.

Biomass was resuspended in lysis buffer (20 mM HEPES-NaOH pH 8.0, 1 M NaCl, 1 mM EDTA, 5% glycerol, 0.1% Tween-20, 2,000 U DNase I, Halt protease inhibitor cocktail, and 1 mM PMSF). Cells were lysed by sonication on ice and clarified by centrifugation (35k × *g*). Clarified lysate was loaded onto a StrepTrap XT Sepharose column (Cytiva) pre-equilibrated with lysis buffer. After washing, AsCas12a was eluted with elution buffer (20 mM HEPES-NaOH pH 8.0, 1 M NaCl, 5 mM desthiobiotin, 5 mM MgCl_2_, and 5% glycerol).

Eluted protein was concentrated to less than 1 mL using a 30-kDa molecular weight cut-off centrifugal concentrator (Millipore). The affinity tag was removed by overnight digestion with SUMO protease at 4 °C with gentle rotation. The protein was subsequently purified by size-exclusion chromatography on a Superdex 200 Increase column (Cytiva) equilibrated in storage buffer (20 mM HEPES-KOH pH 8.0, 150 mM KCl, 5 mM MgCl_2_, and 2 mM DTT, 50% glycerol) and stored at −20 °C.

For cell-based experiments, Cas12a was expressed in *E. coli* BL21(DE3) cells transformed with pET28a-opAsCas12a-NLS ^50^ (Addgene:199605) via electroporation. Cells were grown in TB media supplemented with 0.4% glucose, 25 µg ml−1 chloramphenicol and 100 µg ml−1 ampicillin at 37 °C until OD600 ∼0.6. Protein expression was induced with 0.5 mM IPTG at 18 °C for 14–16 h. Cells were harvested by centrifugation (2,500 g, 10 min, 4 °C) and stored at −80 °C.

Biomass (40 g) was resuspended in lysis buffer (20 mM Tris-HCl pH 8.0, 1500 mM NaCl, 5 mM 2-ME, 5% imidazole) and lysed by sonication at 4 °C. Clarified lysate (35,000 RCF ultracentrifugation) was applied to a HisTrap Ni-NTA column (Cytiva) pre-equilibrated in Buffer A (20 mM Tris-HCl pH 8.0, 500 mM NaCl, 5% imidazole, 5 mM 2-ME) on an ÄKTA Avant system (Cytiva). Nonspecific proteins were removed with Buffer A containing 5% Buffer B (20 mM Tris-HCl pH 8.0, 500 mM NaCl, 500 mM imidazole, 5 mM 2-ME), and target protein was eluted with a linear gradient (5–100% Buffer B). Fractions (3–11) were analyzed by SDS-PAGE.

Pooled fractions were applied to a HiTrap Heparin HP column (Cytiva) pre-equilibrated in Buffer C (20 mM Tris-HCl pH 8.0, 250 mM NaCl, 5 mM 2-ME, 5% glycerol). Protein was eluted with a linear gradient (0–100% Buffer D: 20 mM Tris-HCl pH 8.0, 1000 mM NaCl, 5 mM 2-ME, 5% glycerol). Fractions (11–14) were analyzed by SDS-PAGE.

Final purification was performed by size-exclusion chromatography on a Superdex 200 Increase column (Cytiva) pre-equilibrated in Buffer E (20 mM Tris-HCl pH 8.0, 500 mM NaCl, 5 mM 2-ME). Fractions (13–17) were pooled, dialyzed into storage buffer (20 mM Tris-HCl pH 8.0, 500 mM NaCl, 1 mM DTT, 50% glycerol), and stored at −20 °C.

### *In-vitro* AsCas12a Cleavage Assay

To perform cleavage assays, RNPs were formed by mixing 300 nM of AsCas12a with 900 nM of sgRNA (1:3 concentration) in cleavage buffer (20 mM HEPES, pH 7.5, 150 mM KCl, 10 mM MgCl2, 2 mM DTT) and incubating at room temperature for 15 mins. 10 nM of fully-tagged target DNA was combined with the RNP (100 nM) at 22°C, and cleavage was stopped at different timepoints (0, 10, 30, 90, 270, 810, 2400 seconds) with quenching solution (final concentrations: 0.2U/µl proteinase K (Thermo Fisher Scientific), 0.4M EDTA). Samples were incubated for 30 mins at 37 °C to allow for protein degradation. For the 0 sec timepoint, RNP active mix was stopped with quenching solution and incubated at 37 °C for 30 mins and then DNA target was added.

### Denaturing PAGE gel electrophoresis

Cleaved DNAs were resolved on 15% denaturing PAGE gels (7M urea, large format: 22×18 cm): Each sample was mixed in equal parts with 2X denaturing sample loading buffer (95% formamide, 18 mM EDTA, 0.025% Orange G) and denatured at 90 °C for 3-5 mins followed by cold shock on ice. Each gel was pre-run (200 V, 1 hour, 1X TBE buffer) and then the wells were washed with TBE using a syringe to remove the residual urea from wells. Samples were loaded, and the gel was run in the dark (16 hours, 200 V to eliminate band distortion). The gel was visualized on Amersham Typhoon scanner (Cytiva) by laser excitation at 649 and 550 nm wavelengths (for ATTO-647N/Cy5 and Cy3 labels, respectively).

### Band quantification

Experimental band intensities and lengths from denaturing PAGE were quantified (Image Lab software; Bio-Rad) and normalized to total signal per lane to satisfy mass conservation. Background subtraction was done using rolling disc method adjusted to lane profile. Ladder lanes were marked as standard and each band in ladder was set to its size. A point-to-point (semi-log) regression method was used to fit the ladder to a standard curve. Unknown bands in experimental lanes were fitted to the standard curves of the ladders to accommodate for the irregularities in gel migration. The bands were fitted to the standard curve of the ladder or ladders in the nearest wells.

### Kinetic modelling of Cas12a cleavage and trimming via ODE system

To quantitatively describe the cleavage and trimming activity of Cas12a on a target substrate, we developed a kinetic model based on a system of ordinary differential equations (ODEs). The model considers a substrate *S*(*t*)that is processed into two classes of products: proximal products 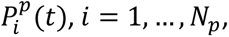 and distal products 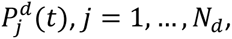 where *N_p_*and *N_d_* denote the number of proximal and distal product species, respectively.

#### Substrate depletion

The substrate is depleted to generate all primary proximal and distal products according to a first-order cleavage reaction under saturating enzyme conditions:

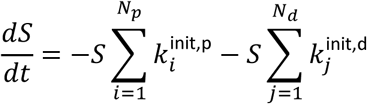

where 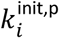 and 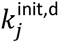 are the rate constants for the formation of the primary proximal *i*-th and distal *j*-th product, respectively.

#### Product dynamics

Each proximal product 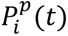 can be formed directly from the substrate and may be trimmed into shorter products. Trimming from a product *i* to product *i* + *d* is governed by a rate constant 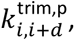 where *d* is the distance between the products in the product series (*e.g* product 1, 2, 3…) The full rate equation for each proximal product is:

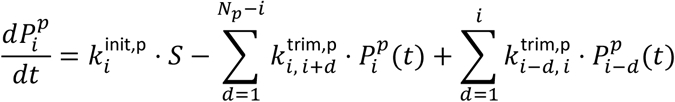

The first term represents formation from substrate, the second term represents loss through trimming into all shorter products, and the third term represents gain from trimming of all larger products. An equivalent formulation applies to the distal product series, with independent rate constants 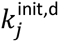 and 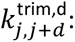

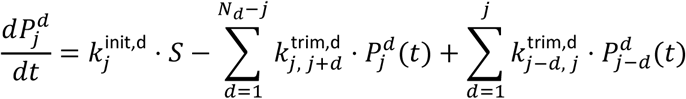

#### Parameter matrix

All formation and trimming rate constants are encoded in a structured rate matrix **K**. Formation rates occupy the diagonal entries and trimming rates at distance *d* occupy the *d*-th super-diagonal, separately for the proximal and distal blocks. This parameterization ensures strictly unidirectional trimming and non-negative rate constants.

#### Model fitting

The system of ODEs was integrated numerically using the LSODA solver (relative tolerance 10^−6^, absolute tolerance 10^−8^) as implemented in scipy.integrate.solve_ivp. Initial conditions were taken directly from the first experimental time point for each species. Rate constants were estimated by minimizing the sum of squared residuals between model predictions and experimental measurements of *S*(*t*), 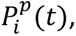 and 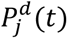 across all time points, using a least-squares optimization routine applied to the flattened rate matrix. Fits that failed to converge were penalized with a large residual (10^6^) to exclude them from the optimum.

### Human cell culture

NK-92 cells (ATCC catalog no.: CRL-2407) were cultured at 37 °C and 5% CO_2_ in modified RPMI 1640 medium (Gibco) supplemented with 12.5% heat-inactivated FBS (Gibco), 1X GlutaMAX-1(Gibco), 1X Penicillin-Streptomycin (Gibco) and 25 mM HEPES (Gibco) and 100 U/ml human IL-2 (PeproTech). They were maintained at 0.8-1 x 10^6^ cells/ml by splitting every 2-3 days to avoid overgrowth.

HEK293T cells (ATCC catalog no.: CRL-3216) were cultured at 37 °C and 5% CO_2_ in DMEM medium with high glucose, GlutaMAX supplement, pyruvate (Gibco) supplemented with 10% heat-inactivated FBS (Gibco) and 1X Penicillin-Streptomycin (Gibco). They were maintained at 70-80% confluency by splitting every 2 days to retain monolayer growth.

SH-SY5Y cells (ATCC catalog no.: CRL-2266) were cultured in 1:1 mixture of Eagle’s Minimum Essential Medium (Minimum Essential Medium (MEM) supplemented with 1x non-essential amino acids and 1mM sodium pyruvate; Gibco) and F12 Medium (Gibco), supplemented with 10% heat-inactivated FBS (Gibco) and 1X Penicillin-Streptomycin (Gibco). The cell cultures were maintained at 37 °C and 5% CO_2_.

### AsCas12a RNP Nucleofection

RNPs were assembled from 33 µM of AsCas12a (with NLS) and 50 µM of gRNA (1:1.5) incubated at 22°C for 20 mins. NK-92 (5 x 10^5^), HEK293T (3 x 10^5^) or SH-SY5Y (2 x 10^5^) cells were counted and spun down to remove media, then washed with 1X PBS (Gibco). The NK-92 cells were resuspended in 18 µl of homemade nucleofection buffer Sol2^51^ (5 mM KCl, 15 mM MgCl_2_, 150 mM sodium phosphate buffer, pH 7.2, 50 mM mannitol) whereas HEK293T and SH-SY5Y cells were resuspended in 18 µl of homemade nucleofection buffer Amaxa V^52^ (5 mM KCl, 10 mM MgCl_2_, 90 mM sodium phosphate buffer, pH 7.2, 20 mM HEPES, 24 mM Sodium Succinate). RNPs (4 µl) were mixed with cells (18 µl) in the corresponding nucleofection buffer and transferred to individual nucleofection cuvette wells (Lonza nucleofection strip-16 well format). Nucleofection was performed in a 4D Nucleofector X-unit (Lonza) with pulse codes CA-137 (NK-92 cells, SH-SY5Y) and FF-120 (HEK293T cells). To facilitate cell recovery, 100 µl of pre-warmed cell culture medium was immediately added to each well and the cells were incubated in the cuvette at 37 °C for 15 mins. Cells were transferred to 24 well plates (Corning) with culture medium and incubated as described (Cell culture method). For SH-SY5Y cell culture, plates were pre-coated with poly-l-lysine (Sigma-Aldrich, 0.001% in PBS) for at least 20 minutes at 37°C.

### Genomic DNA extraction, target gene amplification and nanopore sequencing

After 3 days of growth, NK-92, HEK293T and SH-SY5Y cells were collected by spinning the cells at 1000 x g, followed by a PBS (1X) wash. Genomic DNA was extracted from the cell pellet (Quick-DNA Miniprep Plus Kit; Zymo Research). Target sites were amplified from 100ng of genomic DNA using 30 PCR cycles (Phusion Plus Polymerase, Thermo Fisher Scientific), and then purified (Quick-DNA Miniprep, Zymo Research). Amplified target DNAs were submitted for nanopore sequencing (SeqVision).

### Genome editing analysis pipeline

To analyze nanopore sequencing data, the Linux-based CRISPResso2(v2.3)^25^ pipeline was used in batch mode (for each gene/cell type). A batch file was created in tab-separated (TSV) format (downloaded from CRISPResso2 webpage) where filename for each single end read file (fastq) generated from nanopore sequencing along with reference amplicon and gRNA sequences were indicated. This batch file along with the fastq files were used as input. Default minimum alignment score (homology score to the reference amplicon) was set to 60, and 15 bp were excluded from each side of amplicon (to cover for sequencing start and end errors). INDELs were detected around cleavage site with a quantification window covering 30 base pairs around the gRNA and a cleavage offset of 1 (cleavage position for AsCas12a; default is set for an SpCas9 cleavage site). Top edited reads were selected and used for downstream analysis (gated at >0.9 % reads).

### Statistics and visualization

R-squared values for each individual rate were calculated while modeling AsCas12a cleavage kinetics using our Python-based ODE model implementation. Matplotlib was used for plotting rates for ODEs. Heatmaps and bar graphs were generated using GraphPad Prism. nMDS plots and PERMANOVA analysis were performed using PAST statistical software ^53^. Bray-Curtis similarity index was used to perform PERMANOVA (permutations n=9999) and to generate nMDS plots in 2 dimensions.

## Data and software availability

Raw sequencing data is available through European Nucleotide Archive database with the project accession number PRJEB124505. The ODE system notebooks for processing data from cleavage experiments are available on GitHub (https://github.com/JonesLabEU/gRNA-reprogramming).

## Acknowledgements

We thank Dr. Mindaugas Zaremba for providing AsCas12a for the *in vitro* studies, Dr. Numan Ullah for opAsCas12a-NLS biomass production and Dr. Arunas Silanskas for its purification. We thank Dr. Maria Fernanda Torres Jimenez for help with statistical analysis. We thank all members of Jones lab for their support and help in preparing the manuscript.

SKJ acknowledges funding from the European Research Council, Project 101078247-PROTEGE, and funding from the Research Council of Lithuania under the EMBO Installation Grant program, Project 5826. UA acknowledges funding from the Research Council of Lithuania under the Lithuania-Taiwan Project S-LT-TW-24-12. UN acknowledges funding the Research Council of Lithuania supporting EDV under the competitive PhD project S-PAD-23-11, “The application of gene editing tools for monogenic lysosomal storage disorders”.

## Author Contributions

Conceptualization: UA, SKJ. Methodology: UA, FM, EDV, UN, SKJ. Software: UA, MV, SKJ. Validation: UA, FM. Formal analysis: UA, SKJ. Investigation: UA, FM, EDV. Resources: UN, SJK. Data curation: UA. Writing: UA, SKJ. Editing: UA, FM, MV, EDV, UN, SKJ. Supervision: UA, UN, SKJ. Project administration: SKJ. Funding acquisition: UN, SKJ.

## Supplemental Figures

**Figure S1:**
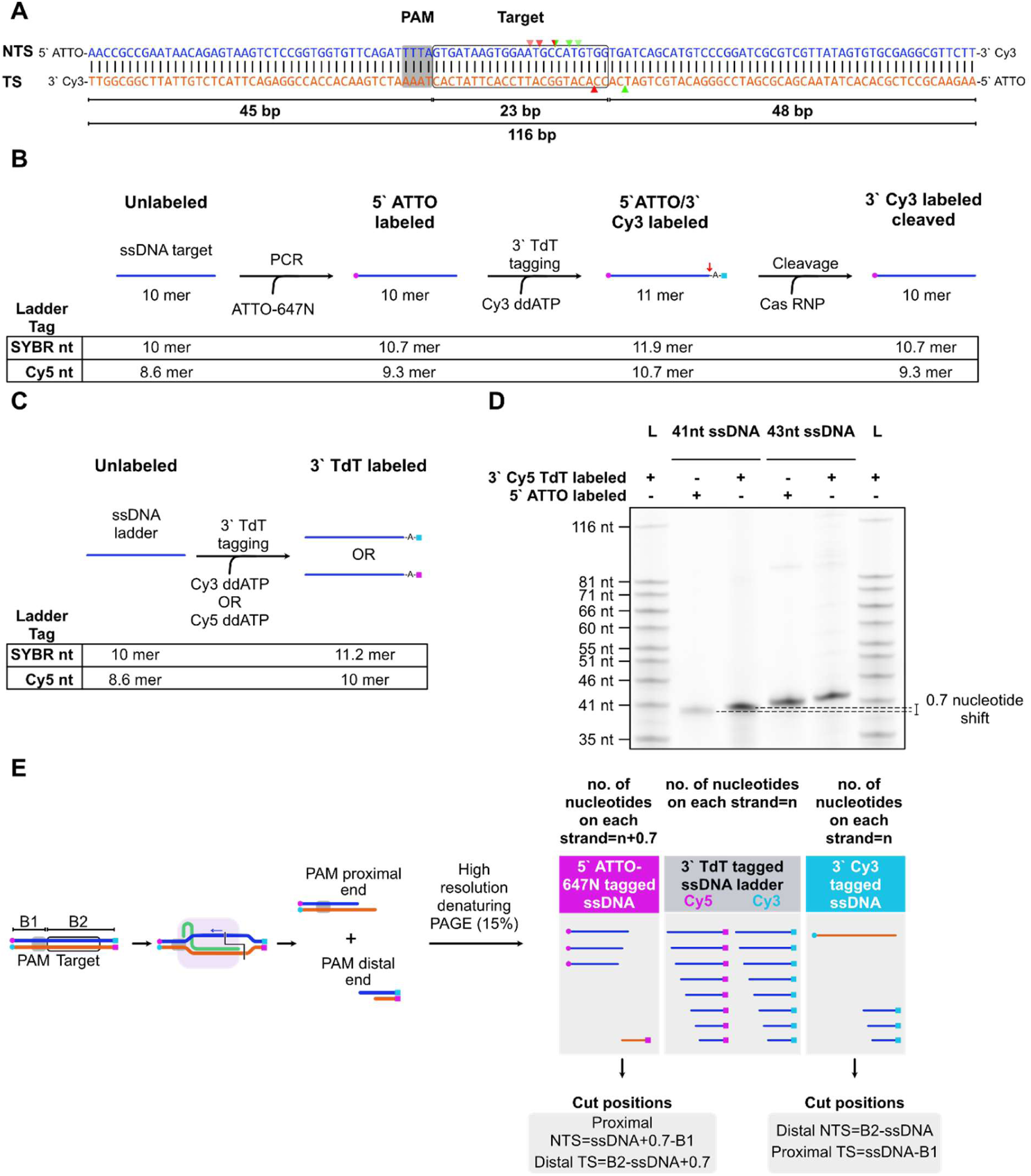
End labelling strategies for DNA targets and cut position calculations. **A)** Schematic of matched target DNA with its cut positions. Red triangles indicate cut positions observed on PAM proximal ends whereas green triangles indicate cut positions observed at PAM distal ends. Triangle with both color shows common cut position observed on both ends. Lighter triangles indicate trimming on NTS. **B)** Schematic of 5’ end labelling with PCR and 3’ end labelling with terminal deoxynucleotidyl transferase (TdT). SYBR nt: The length (in nucleotide equivalents) of an oligonucleotide, compared to an unlabelled one, once stained with SYBR Gold. Cy5 nt: The length (in nucleotide equivalents) of an oligonucleotide, compared to an Cy5 labelled one. **C)** Custom ladder preparation by 3’ TdT labelling with Cy5 and Cy3 ddATP. SYBR nt: The length (in nucleotide equivalents) of an oligonucleotide, compared to an unlabelled one, once stained with SYBR Gold. Cy5 nt: The length (in nucleotide equivalents) of an oligonucleotide, compared to an Cy5 labelled one. **D)** Single nucleotide resolution denaturing urea-PAGE showing differences between 5’ PCR ATTO-labelled DNA and 3’ TdT Cy5 labelled DNA. **E)** Schematic representation of target DNA design and calculations of cut positions by comparing TdT 3’ Cy5 labelled ladder with 5’ ATTO labelled cut products. B1: DNA buffer for proximal end, B2: DNA buffer for distal end.

**Figure S2:**
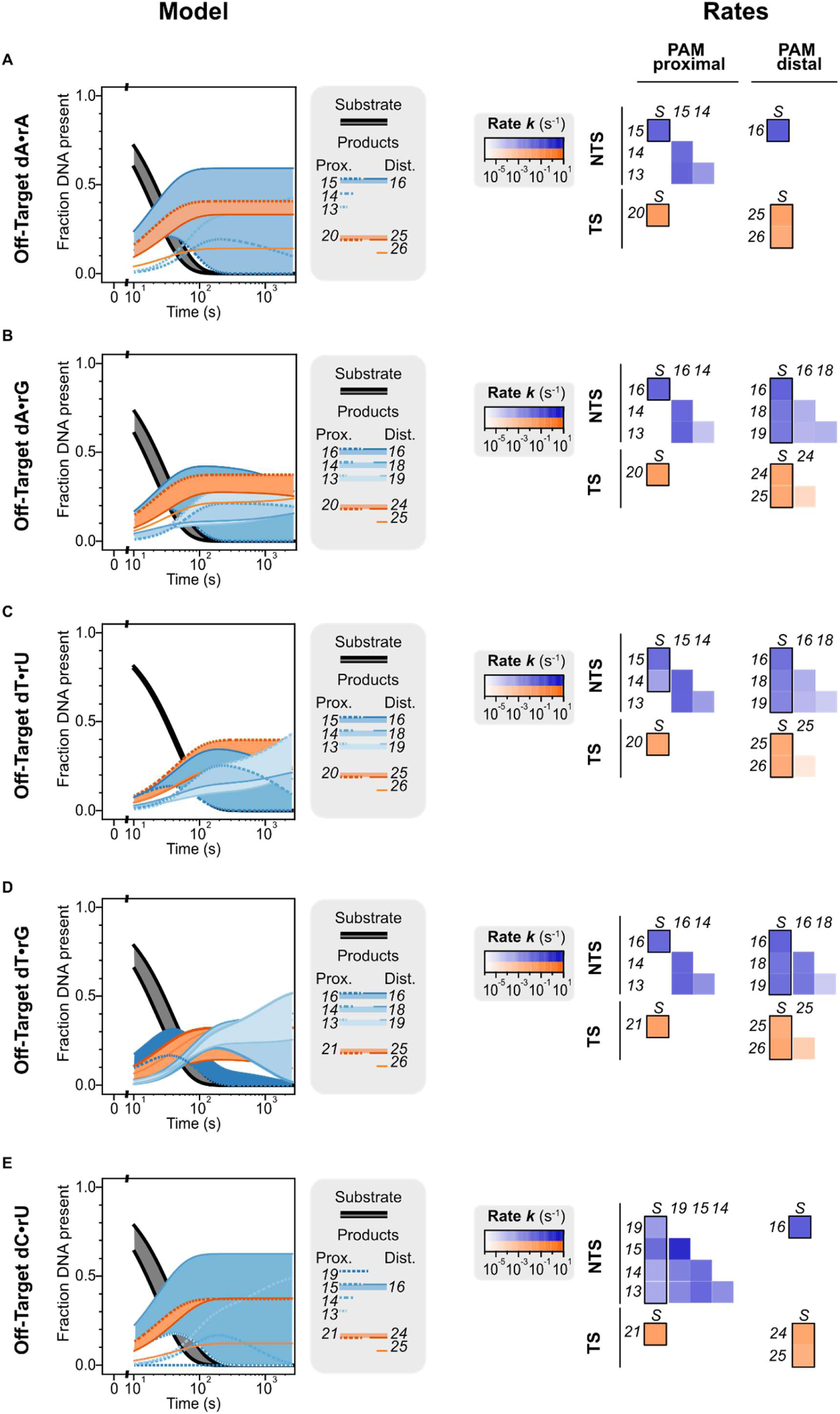
Kinetics of Cas12a cleavage and trimming for different MM19 gRNA and target combinations. **Left:** Complete modelled fractions of Cas12a substrates and products for all DNA ends, where MM19 is dA•rA **(A),** dA•rG **(B),** dT•rU **(C),** dT•rG **(D)** or dC•rU **(E)**. Substrate depletion rates are shown in black, NTS products in shades of blue and TS in orange. Thick lines: PAM-proximal ends, thin lines: PAM-distal ends. Fills assist with identifying product curves. **Right:** Model-fitted rates of conversion from substrates (S) to products, or from one product to another, for the corresponding MM19 gRNA DNA combinations. Numbers identify products by the distance from their cleavage site to the PAM. Blue shades: log-scale rates for the NTS. Orange shades: log-scale rates for the TS. Bolded boxes: products initially cleaved from the substrate.

